# OPAM, an Open source, 3D printed Low-cost Micro-Manipulator for Single Cell Manipulation

**DOI:** 10.1101/2021.08.09.455588

**Authors:** Jiang Xu, Zhuowei Du, Paul Liu, Yi Kou, Lin Chen

**Affiliations:** Molecular and Computational Biology, Department of Biological Sciences, University of Southern California, Los Angeles, CA, 90089, United States; Department of Chemistry, University of Southern California, Los Angeles, CA, 90089, United States; USC Norris Comprehensive Cancer Center, Keck School of Medicine, University of Southern California, Los Angeles, CA 90089, USA

## Abstract

We introduce OPAM, an Open source, low-cost (under $150), 3D-Printed, stepper motor driven, Arduino based, single cell Micromanipulator (OPAM). Modification of a commercial stepper motor led to dramatically increased stability and maneuverability of the motor, based on which the micromanipulator was designed. All components of this micromanipulator can be 3D printed using an entry-level 3D printer and assembled with ease. With this single cell manipulator, successful targeted single cell capture and transfer was confirmed under the microscope, which showed great promise for single cell related experiments.

## Introduction

Recent years have seen a large increment in the needs for single cell related research. Devices either use microfluidics-manufactured through soft lithography [1–6], for high throughput single cell processing, or more complicated devices using electric, magnetic, electromagnetic and mechanical forces to facilitate the manipulation of single cell(s). In some genome structure related research, targeted single cell treatments under the microscope are usually followed by transferring to containers such as PCR tubes for further enzymatic treatments. Such a task is very difficult without special tools. Commercial single cell transferring devices are available but are quite expensive and are not always suitable for special needs. We chose one of the micromanipulation methods--single cell transportation (transfer) [4], which is the task of capturing the desired cell and transferring it to a container such as a PCR tube with minimum volume of liquid (Figure 1a). Such a task seems quite challenging but based on a simple calculation, we found a cheap, yet seemingly precise stepper motor (Figure 1b, left column) may be used to achieve such a goal.

**Figure 1:**
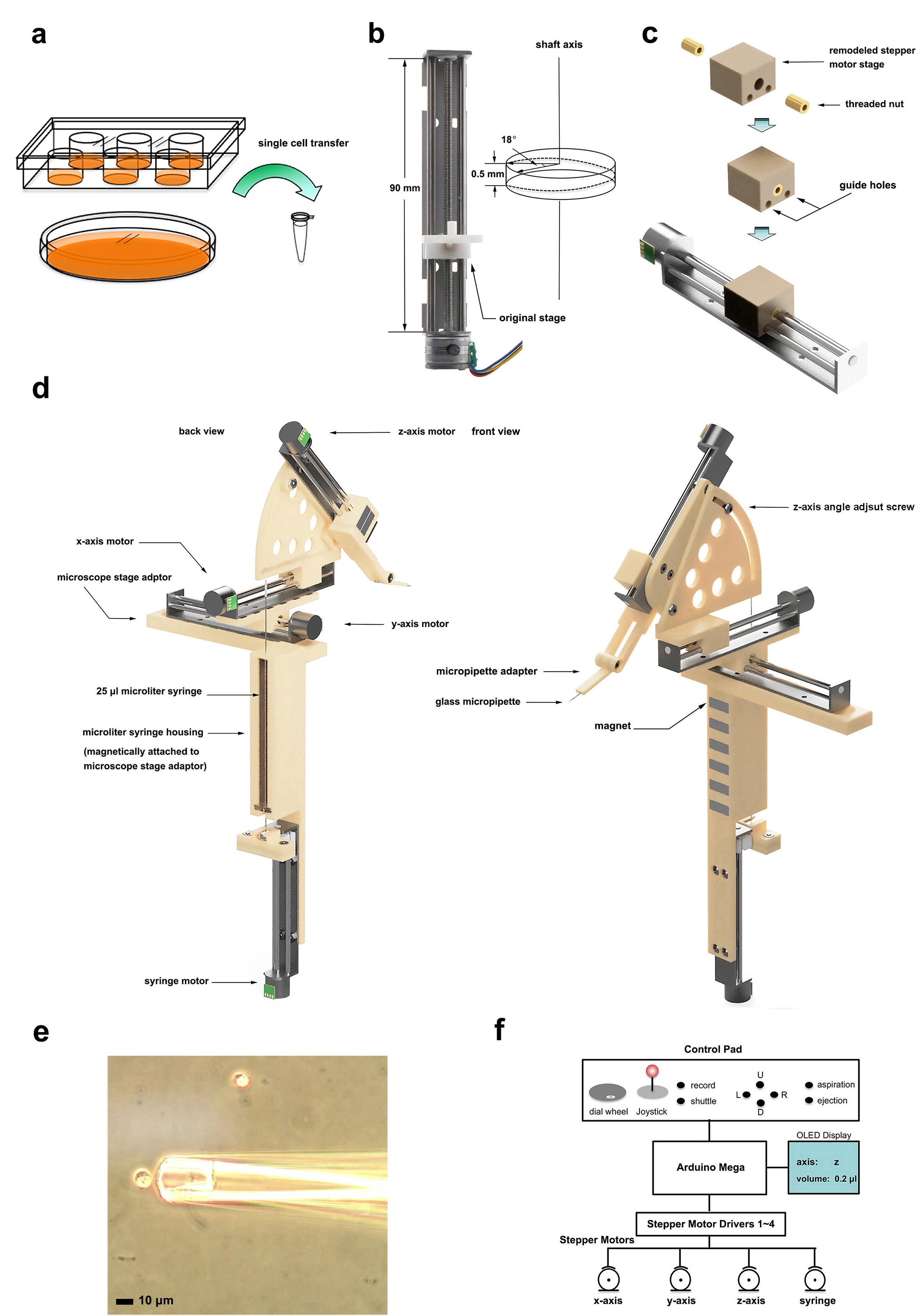
a. The main purpose of OPAM, i.e., transfer of single cell from culture vessels to other containers such as a PCR tube; b. The photo of the low-cost stepper motor used in the current study; c. Remodeling of the moving stage of the stepper motor for increased stability; d. 3D Illustration of the OPAM’s parts and configuration. e. comparison of the size of the GM12878 cells and the micropipette. f. Diagram of the controlling system of OPAM.

This stepper motor was equipped with a 90 mm length screw shaft with a screw pitch of 0.5 mm (Figure 1b, left column), and had a step angle of 18° (Figure 1b, right column). In theory, rotation of such a motor by a single step should move the stage forward (or backward) by 25 μm. With the EasyDriver stepper motor driver, up to ⅛ micro stepping could be enabled, which further decreased the theoretical shortest movement of the stage down to ~ 3 μm. In comparison, the GM12878 cell, a non-attached human lymphoblast cell line commonly used for 3D genome research [7,8], has an average diameter of around 10 μm. From the above calculation it seems that this stepper motor could potentially be used for such a single cell transfer challenge. The biggest challenge is however, not only to move in steady and tiny steps, but also to catch and release the cell with confidence. Potential choices include microprobe, micropipette[9,10], and optical tweezer[11]. Even though optical tweezer is convenient and capable of moving target object(s) under the microscope with ease, the cost of building such system is way much higher, and most importantly, it does not meet our final goal of capturing and transferring the cell of interest out of the original container and to somewhere else, such as a PCR tube for other experiments. The easiest and most affordable option should be a piece of glass micropipette specially crafted for single cell experiment [10,12], which have also been adopted by commercial systems, such as the PicoPipet Pick system (Bulldog Bio) or the UnipicK™ (NeuroInDx, Inc.). However, for more versatile and purpose-oriented applications, an open-source single cell micromanipulator is always desirable. In this article, we describe how we use 3D printing to craft a low cost, robotic micromanipulation system for single cell transfer experiments. We designed a microcontroller-based system to control the moving, aspiration and discharge of the single cell, aiming to make the system reliable and easy to use. Our result showed the great potential of this device for single cell transfer applications, which should be of general interests in the field.

### 3D Design, circuit design and coding

3D design and modeling were done in Fusion 360 (AutoDesk Inc, Educational License) and 3D printing was done with Polylactic acid (PLA) plastic using 3d printer CR-10s (Creality 3D Technology Co., Ltd). As described in the introduction, the low-cost commercial stepper motor could potentially be used for the micromanipulation applications. However, we found that the stage that came with the stepper motor was too loose to allow accurate control. Because there’s only one threaded nut fitted to the screw shaft, and the gap between the stage and guide rails was too wide that the stage was not stable enough for single cell manipulations. To improve the performance of this stepper motor, we redesigned and 3D-printed a new stage with two coaxially placed threaded nuts separated 2 mm away, and with tighter guide holes for the stage to slide on the rails (Figure 1c). The two guide holes were polished with a drill press coupled with a 3 mm drill bit, which was the same as the rail shaft. With such modifications, the stage became much more stable, and moved more smoothly along the rail. To attach the micromanipulator to our inverted microscope (Olympus, CKX41), we designed, and 3D printed a piece of magnetic adapter and screwed it onto the stage of the microscope (Figure 1d). The micromanipulator was based on a modular design that used magnets to hold each module together. Based on this notion, the z-axis assembly, including the z-axis stepper motor and the sector shaped rack was stacked onto the moving stage of the x-axis stepper motor. The x-axis stepper motor was stacked onto the moving stage of the y-axis stepper motor, which was stacked onto the microscope adaptor. The z-axis stepper motor was attached to a sector shaped rack, enabling the angle between the z-axis and the x-y plane to be adjustable. The micropipette adapter was attached to the moving stage of the z-axis via two pieces of magnets. The microliter syringe housing was also attached to the microscope adaptor via one piece of magnet and is adjustable (Figure 1d). Such modular design makes assembly and attachment / detachment of the micromanipulator onto the stage of the microscope much easier while providing the stability and versatility needed for single cell manipulations. Finally, the microliter syringe was connected to the glass micropipette via a piece of silicon capillary tube (not shown in the diagram in Figure 1d).

For digitalized control of the four axes (x, y, z, and syringe), an Arduino Mega microcontroller was used to control the four EasyDriver stepper motor drivers (SparkFun Electronics), which in turn drive the four stepper motors respectively (Figure 1f). A Joystick with built-in click switch was used for gross movement of the motors of the x, y and z and toggle between the axes (The x-y plane and z axis). Four push button switches (up, down, left, right) were used for additional stepwise control of the stepper motors. A dial wheel switch (rotary encoder) was used for selecting the desired number of steps for the selected stepper motor to move. Clicking the dial wheel will make the stepper motor to move the desired micro steps. Two additional push button switches were used for controlling the microliter syringe for fast aspiration or ejection (Figure 1f). To make the operation easier, we implemented another two push button switches, one to record the route and the other to shuttle back and forth between the source location and the target location automatically. Pressing the ‘record’ button would record the position of the micropipette. The current version of the control code is capable of recording dozens of such positions. In practice, one needs to manually (by pressing the buttons) move the micropipette from the target location to the source location and to press the ‘record’ button to save the coordinate information when the micropipette stops at a key position. Pressing the shuttle button would shuffle the micropipette between the first coordinate and the last coordinate saved, which would be quite convenient for transferring multiple single cells from the cell suspension to PCR tubes.

For monitoring the status of the device, a 0.96” OLED display was used for displaying key information such as coordinate, micro steps, and position information. The code for the micromanipulator was written in C++ in the Arduino IDE and loaded to the Arduino Mega. The detailed circuit diagram was shown in supplemented Figure 2.

## Results

### Modified stepper motor stage showed much higher stability upon which the micromanipulator was built

The precision of the entry-level 3D printing was just enough to make the two threaded nuts align well. After reinstalling the custom designed stage, the movement was much more stable, at the expense of increased resistance. To solve the problem, we applied grease to the threaded shaft and rail shaft and increased the driving voltage from 6 volts to 12 volts. The stage was then freely movable in both directions. The whole system was based on this modified version of the stepper motor and was assembled using the 3D printed parts, magnets and 3 mm bolts and nuts (Figure 1d).

### Microliter syringe coupled with the motorized housing enabled precise control of fluid flow and single cell transfer with custom pulled micro pipette

To produce the glass micropipette, we pulled the glass capillaries (3.5″ Drummond, Item# 3-000-203-G/X) with a capillary puller (P-2000 Laser-Based Micropipette Puller, Sutter Instrument company). The resultant micropipette had a very fine tip that need to be trimmed according to the size of the cells. Under the microscope, we trimmed the tapered ends of the micropipette with a blade to get an orifice with diameter like the cell (Figure 1e). To gain more accurate control of the micro-pipetting process, we designed and crafted a motorized housing for the microliter syringe (Hamilton) using the same, but unmodified stepper motor mentioned above (Figure 1d). We managed to get several usable glass micropipettes that had suitable bore size. We then inserted the micropipette to the micropipette adaptor (Figure 1d), filled the microliter syringe with PBS buffer and connected the micropipette to the needle of the microliter syringe via a piece of silicon capillary (not shown in Figure 1d). The syringe was then controlled to expel the trapped air bubble inside the micropipette until the buffer formed a droplet at the tip of the micropipette, and the system priming was finished and ready to use.

For testing single cell transferring, 1% formaldehyde pre-fixed GM12878 cells [<u>7,8,12</u>] were diluted at 4×10^5^/ml in PBS buffer. 10 μl of such cell suspension was transferred to a piece of glass slide, which was then mounted onto the stage of the inverted microscope. We then controlled the micropipette to move to a nearby location and slowly approached the cell of interest, rotated the dial wheel to select the desired number of steps of the syringe motor and clicked the dial wheel to aspirate the cell into the micropipette. We then controlled the micromanipulator to move the micropipette to another location, selected the desired number of steps of the syringe motor and clicked the dial wheel to discharge the cell. The result was very encouraging that we could repeatedly pick up a desired single cell at one location, move to another location under the microscope and discharge the cell with confidence (See Supplemented Video). Generally, we could transfer the cell using less than 0.5 μl of liquid, which is compliant with many downstream experiments.

## Discussion

Using four pieces of low-cost commercial stepper motors, a microliter syringe, an Arduino Mega microcontroller, four EasyDriver stepper motor drivers, and custom designed and 3D printed parts we built an open source, digitally controlled micromanipulator that could transfer single cell(s) to desired locations. The use of an Arduino-based controller made the design of the control system much easier for non-professional amateur engineers. Entry level 3D printing, on the other hand, made the crafting of the whole system with ease and provided a certain level of precision. The result, however, was far beyond our expectation that the remodeled 3D printed stepper motor stage provided much higher stability, capable of doing more precise work than the original one. Due to precision limitation in measurement and crafting processes, the revised stage had higher resistance with the two guide rails that we had to increase the driving voltage from 6 volt to 12 volt to compensate. With better measurement and tools, the precision of the part should be increased, and the stepper motor should be controlled more sensitively. We also found that the micromanipulator’s reaction was a bit sluggish when changing the moving directions (i.e., from left to right, or from up and down) at each axis, which should be attributed to the native gap between the screw thread of the shaft and nut (Supplemented Figure 1a). Such a problem can be overcome through forcing the threads of the threaded nut against the thread of the shaft in both directions with a pair of springs, which can minimize the delays when changing moving directions (Supplemented Figure 1b).

We tested our micromanipulator for the transferring of formalin fixed GM12878 cells under the microscope, from which we could clearly see whether the transfers were successful or not. From multiple rounds of tests, we were able to demonstrate that single cell being captured into the micropipette, transferred to another location, and discharged, which could be alternatively discharged into a PCR tube. The only problem with the micropipette approach is that for some micropipettes, even though the cell can be captured and released, the discharged cell sometimes would stick to the orifice of the micropipette, making the transfer result unknown if we should transfer the cell to a PCR tube without visual confirmation. This might be caused by the rough surface of the orifice resulted from blade cutting. Ideally the orifice should be smooth and perpendicular to the longitudinal axis of the micropipette. In practice, we found it’s hard to obtain such an ideal micropipette with the blade cutting approach. All the orifices we got from blade cutting are of irregular shapes, which made some micropipette not being able to pick up the cell or having the cell adhesion problem during discharging of the cell. Also, the wall of the glass capillaries we used to produce the micropipette was too thick, making the micropipette bore surface area too large, which might be another factor leading to the cell adhesion problem.

The current design, albeit still has many problems, has shown great potential of these low-cost stepper motors of being modified and assembled as the robotic arm to carry out meticulous work at single cell level. With improvement addressing the problem mentioned above, the performance of this low-cost single cell manipulator should be further improved. Because of its small size, our micromanipulator can be easily attached to many microscopes, from simple inverted microscopes in the cell culture room to more advanced fluorescence microscopes, enabling transfer of cells of interest with ease. The digital control system enables precise and remote control and thus can be placed anywhere in the lab where human accesses are restricted. We have released all the information, including 3D designs, coding and circuit diagram to our GitHub page and will update new versions in the future. Anyone who are interested in single cell transfer or micromanipulation can make modifications to meet their own purposes.

## Contributions

JX conceived the idea, designed, crafted the model, made the circuit, wrote the code, and tested the single cell transfer experiment. ZWD pulled the glass capillary, JX produced the glass micropipette, PL and YK did the cell culturing.

## Design Availability

3D model, code and circuit diagram can be found at https://github.com/JiangXu123/DIY_Lab_Instrument/tree/master/micro_manipulator

## Acknowledgement

We thank all the friendly folks from the Arduino forum (https://forum.arduino.cc/) for providing valuable suggestions and discussions.

---

Supplemented Figure 1, a. Illustration of the principle of how sluggish movement occur during change of moving directions from moving left to moving right. b. Illustration of how two springs could solve the sluggish movement during change of directions.

Supplemented Figure 2, Circuit diagram of OPAM (Drew with EasyEDA)

Supplemented Video, 00:00~00:03: Experimental setup with the most primitive version of OPAM for single cell transfer experiment; 00:04~04:18: Four clips (clip1 to clip 4) showing successful capture, transfer, and discharge of single cell; 04:18~end: Seven clips (clip 5 to clip 11) showing successful capture, transfer, and discharge of single cell, with cell stick to the orifice of the micropipette after discharge.

