## Supplementary figures and images for "OPAM, an Open source, 3D printed Low-cost Micro-Manipulator for Single Cell Manipulation"

### Supplemental Figure 1

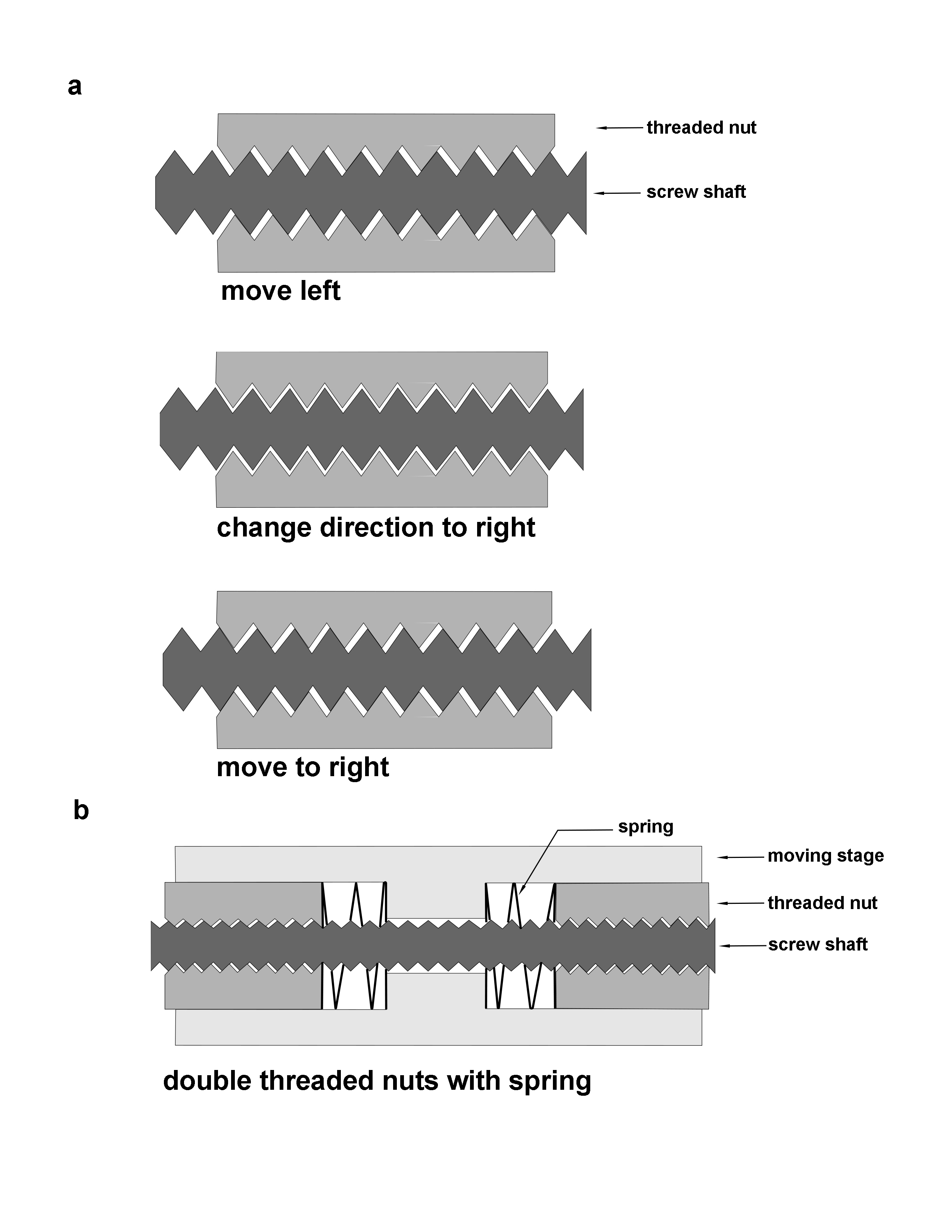

### Supplemental Figure 2

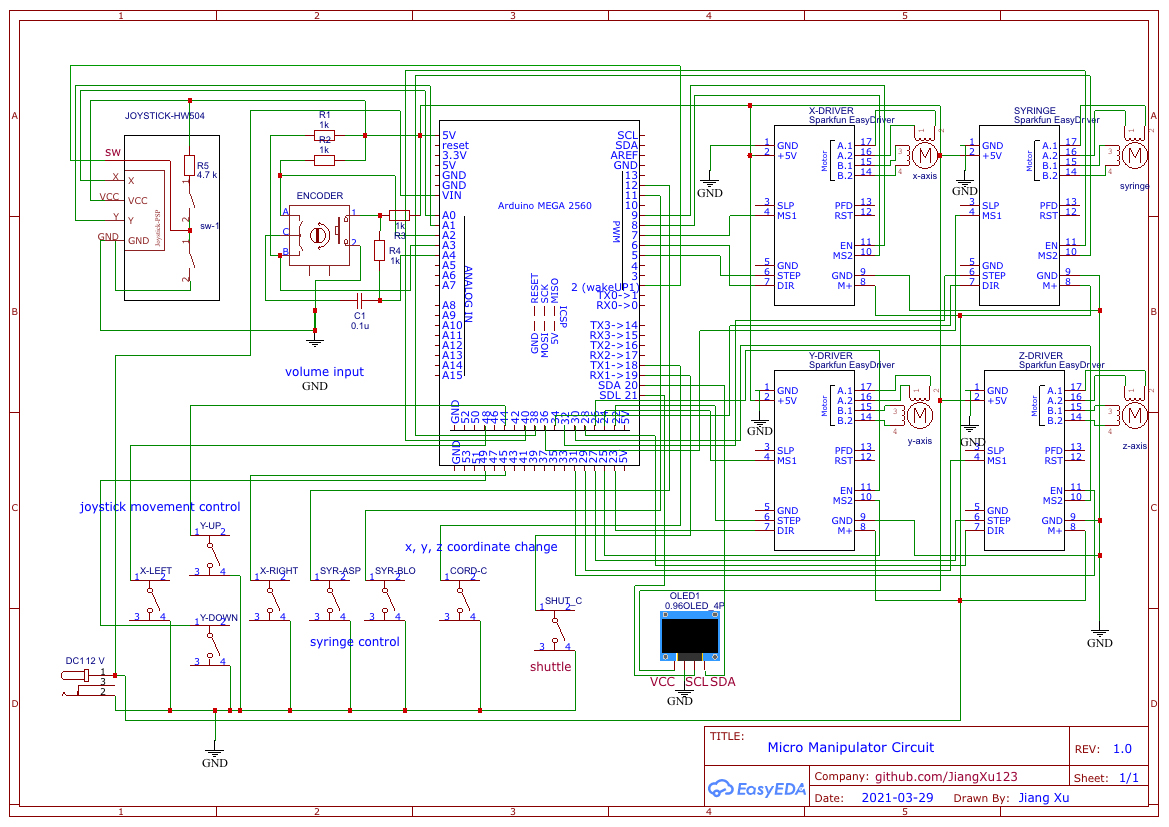
